# Negligible impact of SARS-CoV-2 variants on CD4^+^ and CD8^+^ T cell reactivity in COVID-19 exposed donors and vaccinees

**DOI:** 10.1101/2021.02.27.433180

**Authors:** Alison Tarke, John Sidney, Nils Methot, Yun Zhang, Jennifer M. Dan, Benjamin Goodwin, Paul Rubiro, Aaron Sutherland, Ricardo da Silva Antunes, April Frazier, Stephen A. Rawlings, Davey M. Smith, Bjoern Peters, Richard H. Scheuermann, Daniela Weiskopf, Shane Crotty, Alba Grifoni, Alessandro Sette

## Abstract

The emergence of SARS-CoV-2 variants highlighted the need to better understand adaptive immune responses to this virus. It is important to address whether also CD4+ and CD8+ T cell responses are affected, because of the role they play in disease resolution and modulation of COVID-19 disease severity. Here we performed a comprehensive analysis of SARS-CoV-2-specific CD4+ and CD8+ T cell responses from COVID-19 convalescent subjects recognizing the ancestral strain, compared to variant lineages B.1.1.7, B.1.351, P.1, and CAL.20C as well as recipients of the Moderna (mRNA-1273) or Pfizer/BioNTech (BNT162b2) COVID-19 vaccines. Similarly, we demonstrate that the sequences of the vast majority of SARS-CoV-2 T cell epitopes are not affected by the mutations found in the variants analyzed. Overall, the results demonstrate that CD4+ and CD8+ T cell responses in convalescent COVID-19 subjects or COVID-19 mRNA vaccinees are not substantially affected by mutations found in the SARS-CoV-2 variants.

## INTRODUCTION

The emergence of several SARS-CoV-2 variants of concern (VOC) with multiple amino acid replacements has implications for the future control of the COVID-19 pandemic (Davies et al., 2020; Kirby, 2021; Tegally et al., 2020; Volz et al., 2021). Variants of concern include the UK (United Kingdom) variant 501Y.V1 lineage B.1.1.7 (Davies et al., 2020), the SA (South Africa) variant 501Y.V2 lineage B.1.351 (Tegally et al., 2020), the BR (Brazilian) variant 501Y.V3 lineage P.1 (Voloch et al., 2020) and the CA (California) variant CAL.20C lineage B.1.427 (Zhang et al., 2021). The B.1.1.7 variant is associated with increased transmissibility (Rambaut et al., 2020; Washington et al., 2021), and similar epidemiological observations have been reported for the SA and BR variants (Tegally et al., 2020; Voloch et al., 2020).

Mutations of greatest concern are present in the viral Spike (S) protein, and include notable mutations in the receptor binding domain (RBD), N-terminal domain (NTD), and furin cleavage site region. Several of these mutations directly affect ACE2 receptor binding affinity, which may impact infectivity, viral load, or transmissibility (Greaney et al., 2021; Starr et al., 2021; Wang et al., 2021a; Zahradník et al., 2021). Several of the mutations were also noted to be in regions bound by neutralizing antibodies, so it is crucial to address to what extent the mutations associated with the variants impact immunity induced by either SARS-CoV-2 infection or vaccination.

Several reports address the effect of these mutations on antibody binding and function, by either monoclonal or polyclonal antibody responses, and considering both natural infection or vaccination (Edara et al., 2021; Greaney et al., 2021; Muik et al., 2021; Shen et al., 2021; Skelly et al., 2020; Stamatatos et al., 2021; Supasa et al., 2021; Wang et al., 2021a; Wang et al., 2021b; Wibmer et al., 2021; Wu et al., 2021). In general, the impact of the B.1.1.7 variant mutations on neutralizing antibody titers is moderate (Emary et al., 2021; Muik et al., 2021; Shen et al., 2021; Skelly et al., 2020; Supasa et al., 2021; Wu et al., 2021). In contrast, the mutations associated with the B.1.351 and P.1. variants are associated with more pronounced loss of neutralizing capacity (Cele et al., 2021; Skelly et al., 2020; Wang et al., 2021a; Wibmer et al., 2021; Wu et al., 2021). Concerning vaccination responses, the AstraZeneca ChAdOx1 vaccine has been associated with a partial loss of neutralizing antibody activity against B.1.1.7 (Skelly et al., 2020), and a large loss of neutralizing activity against B.1.351 (Voysey et al., 2021). Consistent with these reports, ChAdOx1 maintains efficacy against B.1.1.7 (Emary et al., 2021; Hall et al., 2021), but has a major loss in efficacy against mild COVID-19 with the B.1.351 variant (Voysey et al., 2021). Current epidemiological evidence is that the BNT162b2 Pfizer/BioNTech COVID-19 vaccine retains its efficacy against B.1.1.7 in the UK and in reports from Israel (Amit et al., 2021). Novavax (NVX-CoV2373) has reported differential protective immunity against the parental strain, B.1.1.7, and B.1.351 in vaccine clinical trials (96%, 86%, and 60%) (Novavax Inc., 2021), whereas the Janssen Ad26.COV2.S 1-dose COVID-19 vaccine, which elicits lower neutralizing antibody titers (Sadoff et al., 2021), has relatively similar protection for moderate COVID-19 against both the ancestral strain and B.1.351 (72% and 64%)(FDA, 2021a, b).

Several lines of evidence suggest that CD4^+^ and CD8^+^ T cell responses play important roles in resolution of SARS-CoV-2 infection and COVID-19 (Sette and Crotty, 2021), including modulating disease severity in humans (Rydyznski Moderbacher et al., 2020; Tan et al., 2021) and reducing viral loads in non-human primates (Munoz-Fontela et al., 2020). Further, persons with agammaglobulinemia or pharmaceutical depletion of B cells generally experience an uncomplicated COVID-19 disease course (Sette and Crotty, 2021; Soresina et al., 2020). Robust CD4^+^ and CD8^+^ T cell memory is induced after COVID-19 (Breton et al., 2021; Dan et al., 2021; Peng et al., 2020; Wang et al., 2021b), and multiple COVID-19 vaccines elicit CD4^+^ and CD8^+^ T cell responses (Baden et al., 2021; Dowd et al., 2020; Keech et al., 2020; Sadoff et al., 2021; Voysey et al., 2021). It is therefore key to address the potential impact of SARS-CoV-2 variants mutations on T cell reactivity; however, little data is currently available on this topic (Skelly et al., 2020).

Here, we take a combined experimental and bioinformatics approach to address how SARS-CoV-2 variants of concern impact T cell reactivity. We directly assess T cell responses from persons recovered from COVID-19 obtained before the emergence of the variants, and from recent Moderna mRNA-1273 or Pfizer/BioNTech BNT162b2 vaccinees, for their capacity to recognize peptides derived from the ancestral reference sequence and the B.1.1.7, B1.351, P.1 and the CAL.20C variants. Bioinformatic analyses were used to predicted the impact of mutations in the various variants with sets of previously reported CD4^+^ and CD8^+^ T cell epitopes derived from the ancestral reference sequence (Tarke et al., 2021).

## RESULTS

### Sequence analysis, peptide pool generation and selection of cohorts of COVID-19 convalescent and recent vaccinees

As a first step, we mapped the specific mutations (amino acid replacements and deletions) associated with several of the current variants of concern, including the SARS-CoV-2 B.1.1.7, B1.351, P.1 and the CAL.20C variants, as compared to the SARS-CoV-2 Wuhan ancestral sequence (NCBI acc. no. NC_045512.2). Briefly, the genomic sequences were downloaded from GISAID, translated using the VIGOR4 tool available on the Virus Pathogen Resource (ViPR) (Pickett et al., 2012), and then compared with the protein sequence of the Wuhan ancestral strain to identify all the possible amino acid changes, as listed in **Table S1**.

Next, we synthesized the corresponding peptides associated with the different variants and generated new peptide pools spanning the full genome sequences of the ancestral Wuhan strain and the respective B.1.1.7, B1.351, P.1 and the CAL20C variants (**Table S2**). As described below, the resulting peptide pools were assessed for their capacity to be recognized by memory T cells responses derived from natural infection in convalescents and vaccinees, and responses to the variant and ancestral genome antigen-specific pools were compared.

Our convalescent donors were adults with ages ranging from 21 to 57 years of age (median 39); 27% were male and 73% female (**Table 1**). SARS-CoV-2 infection in these donors was determined by PCR-based testing during the acute phase of their infection, if available (55% of the cases), and/or seropositivity determined by plasma SARS-CoV-2 S protein RBD IgG ELISA (Stadlbauer et al., 2020)(**Fig. S1**). From these donors, PBMC samples were collected between July to October 2020 period, when the dominant local strain was the ancestral reference virus.

**Table 1.**
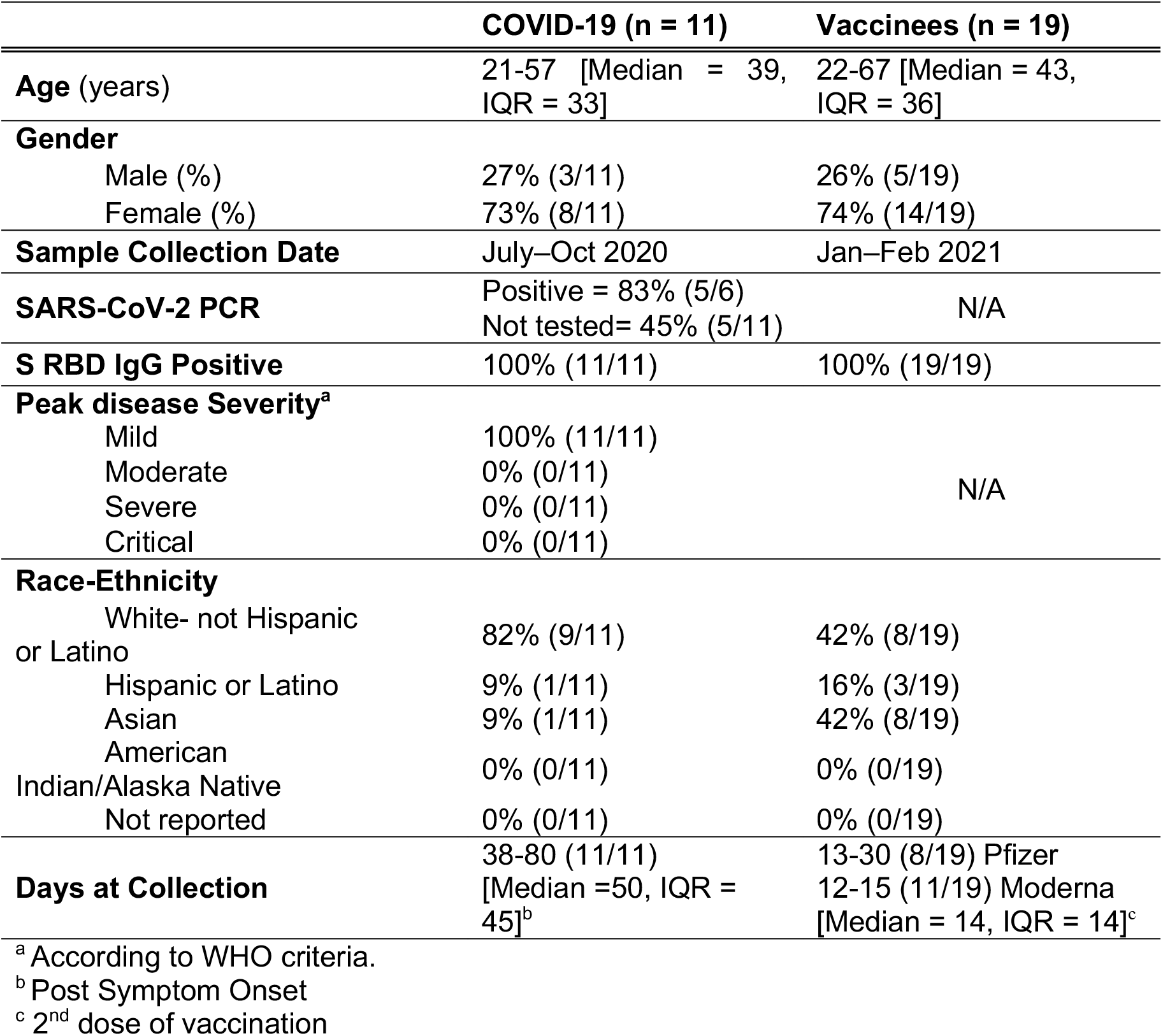
Characteristics of donor cohorts.

From vaccinated donors, we collected PBMC after recent vaccination with the Moderna mRNA-1273 or the Pfizer/BioNTech BNT162b2 vaccines, approximately 14 days following the second dose administration (**Table 1**). These donors ranged in age from 22 to 67 years (median 43) and 26% were male and 74% were female. All vaccinees had significant RBD IgG titers in the 1843 to 16365 range, consistent with recent vaccination **(Fig. S1**).

### CD4^+^ and CD8^+^ T cell antigenicity against Spike variant sequences in convalescent samples

We previously described the use of Activation Induced Marker (AIM) assays (Dan et al., 2021; Grifoni et al., 2020b; Mateus et al., 2020; Rydyznski Moderbacher et al., 2020; Tarke et al., 2021) to measure CD4^+^ and CD8^+^ T cell responses to pools of overlapping peptides spanning the entire sequence of the SARS-CoV-2 antigens. Here, we utilized the same AIM assays using OX40^+^CD137^+^ and CD69^+^CD137^+^ markers for CD4^+^ and CD8^+^ T cells reactivity, respectively (Grifoni et al., 2020b; Mateus et al., 2020). As shown in **Fig. 1A-B**, good CD4^+^ and CD8^+^ T cell reactivity was observed in convalescent donors with pools of overlapping peptides spanning the S protein of the ancestral Wuhan sequence, but also for each of the corresponding variant S pools. Geomean reactivity ranged from 0.09 to the 0.10 for CD4^+^ T cells, and 0.08 to the 0.11 for CD8^+^ T cells; No significant difference was observed between the pool of S peptides corresponding to the ancestral sequence and those corresponding to the different variants (CD4: UK p=0.90; SA p=0.50; BR p=0.49; CA p=0.85 and CD8: UK p=0.16; SA p=0.07; BR p=0.18; CA p=0.20 by the Wilcoxon test). These values (here and in subsequent graphs) are not corrected for multiple comparisons, as the correction would only decrease the statistical power for detecting a significant difference; therefore, not performing multiple comparison corrections is the more conservative and stringent test.

**Figure 1.**
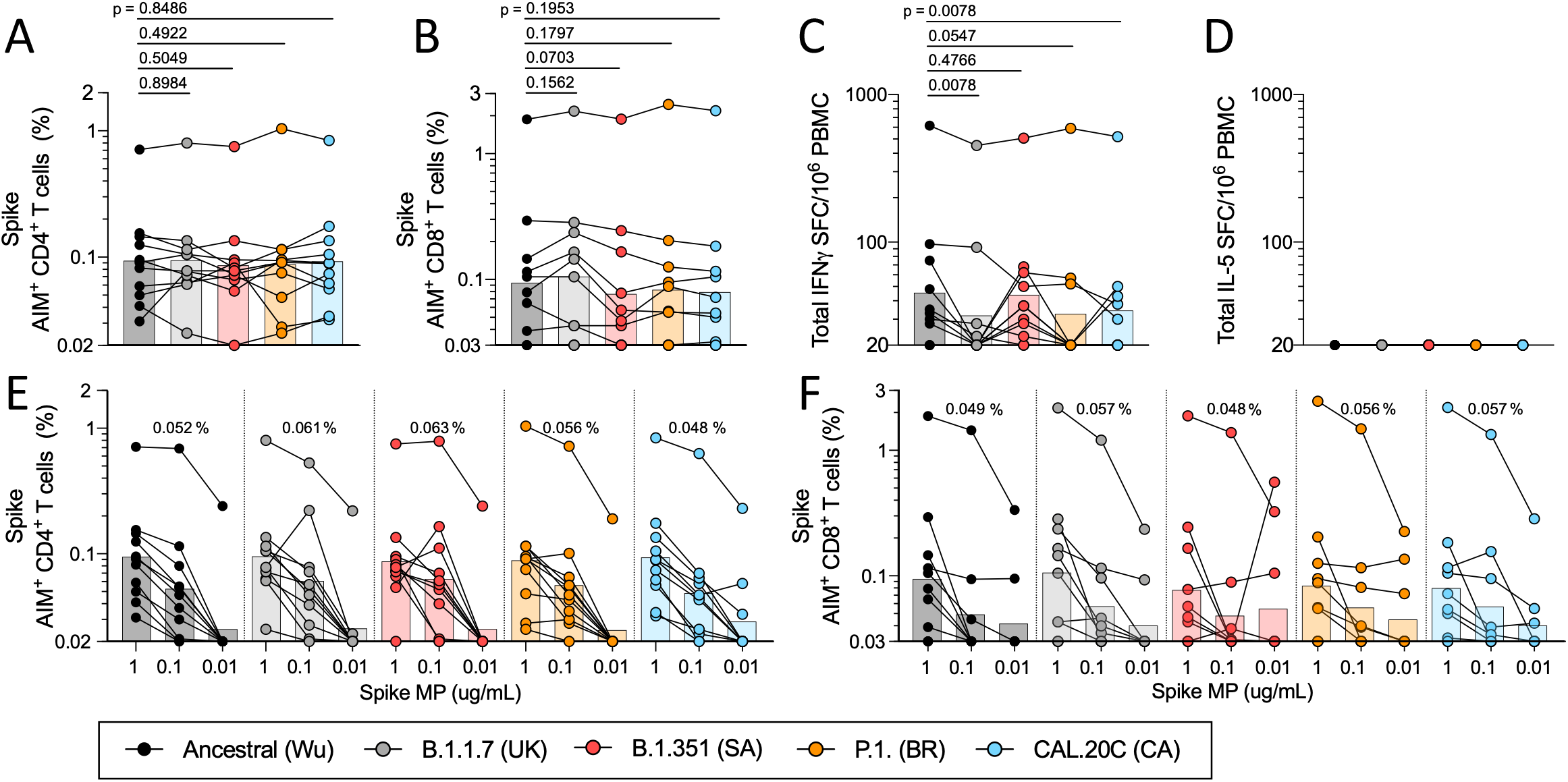
T cell responses of COVID-19 convalescent individuals against SARS-CoV-2 Spike for the different variants. PBMCs of COVID-19 convalescent individuals (n=11) were stimulated with the Spike MPs corresponding to the ancestral reference strain (Wu, black) and the B.1.1.7 (UK, grey), B.1.351 (SA, red), P.1 (BR, orange) and CAL.20C (CA, light blue) SARS-CoV-2 variants. **A**) Percentages of AIM^+^ (OX40^+^CD137^+^) CD4^+^T cells. **B**) Percentages of AIM^+^ (CD69^+^CD137^+^) CD8^+^ T cells. **C**) IFNγ spot forming cells (SFC) per million PBMCs **D**) IL-5 SFC per million PBMCs. Paired comparisons of Wuhan S MP versus each of the variants were performed by Wilcoxon test and are indicated by the p values in panels **A**-**C**. The data shown in panels **A** and **B** are plotted to show the Spike MPs titration (1 μg/mL, 0.1 μg/mL, 0.01 μg/mL) for CD4^+^ (**E**) and CD8^+^ (**F**) T cells in each SARS-CoV-2 variant and the geometric mean of the 0.1ug/mL condition is listed above each titration. In all panels, the bars represent the geometric mean.

These T cell analyses were extended using a FluoroSPOT assay system, to measure the capacity of the various pools to elicit functional responses in terms of secretion of IFNγ and IL-5 cytokines (**Fig. 1C-D)**. As shown in **Fig. 1C**, reactivity was observed for the S pools in convalescent donors in terms of IFNγ with geomean reactivity ranging from 32 to 45 Spot Forming Cells (SFC) per million PBMCs. Compared to the ancestral strain, mild decreases in the 24–30% range were noted for B.1.1.7, P.1 and CAL.20C variant pools (UK p=0.01; SA p=0.48; BR p=0.05 and CA p=0.01 by the Wilcoxon test), while no difference was observed for B.1.351. As expected, no IL-5 reactivity was observed for any of the pools (**Fig. 1D).**

To further expand these findings, we considered the dose response of the various S pools in terms of stimulation of CD4^+^ and CD8^+^ T cell specific responses. As shown in **Fig. 1E**, CD4^+^ T cell dose dependent responses for the Wuhan and four variant pools were similar. The same pattern was also observed for CD8^+^ T cell responses (**Fig. 1F**).

### CD4^+^ and CD8^+^ T cell antigenicity of proteome-wide SARS-CoV-2 variant sequences in convalescent samples

As shown in **Table S1**, mutations found in the variants studied herein were not limited to the Spike protein, but occurred in several additional antigens encoded in the SARS-CoV-2 genome. To address their potential impact on the overall proteome-wide CD4^+^ and CD8^+^ T cell reactivity, we tested overlapping peptide pools spanning the entire proteome of the ancestral Wuhan sequence in comparison with corresponding pools representing the different variants.

Overall, reactivity to the peptide pools spanning the variant genomes was found to be similar to that against the ancestral Wuhan strain (**Fig. 2**). When the sum total of reactivity throughout the genome was considered, no differences or decreases in reactivity compared to the ancestral were noted for the variant pools (CD4: UK p=0.58; SA p= 0.46; BR p= 0.27; CA p= 0.08 and CD8: UK p= 0.25; SA p= 0.15; BR p= 0.02; CA p= 0.30 by the Wilcoxon test uncorrected p values) (**Fig. 2A-B**).

**Figure 2.**
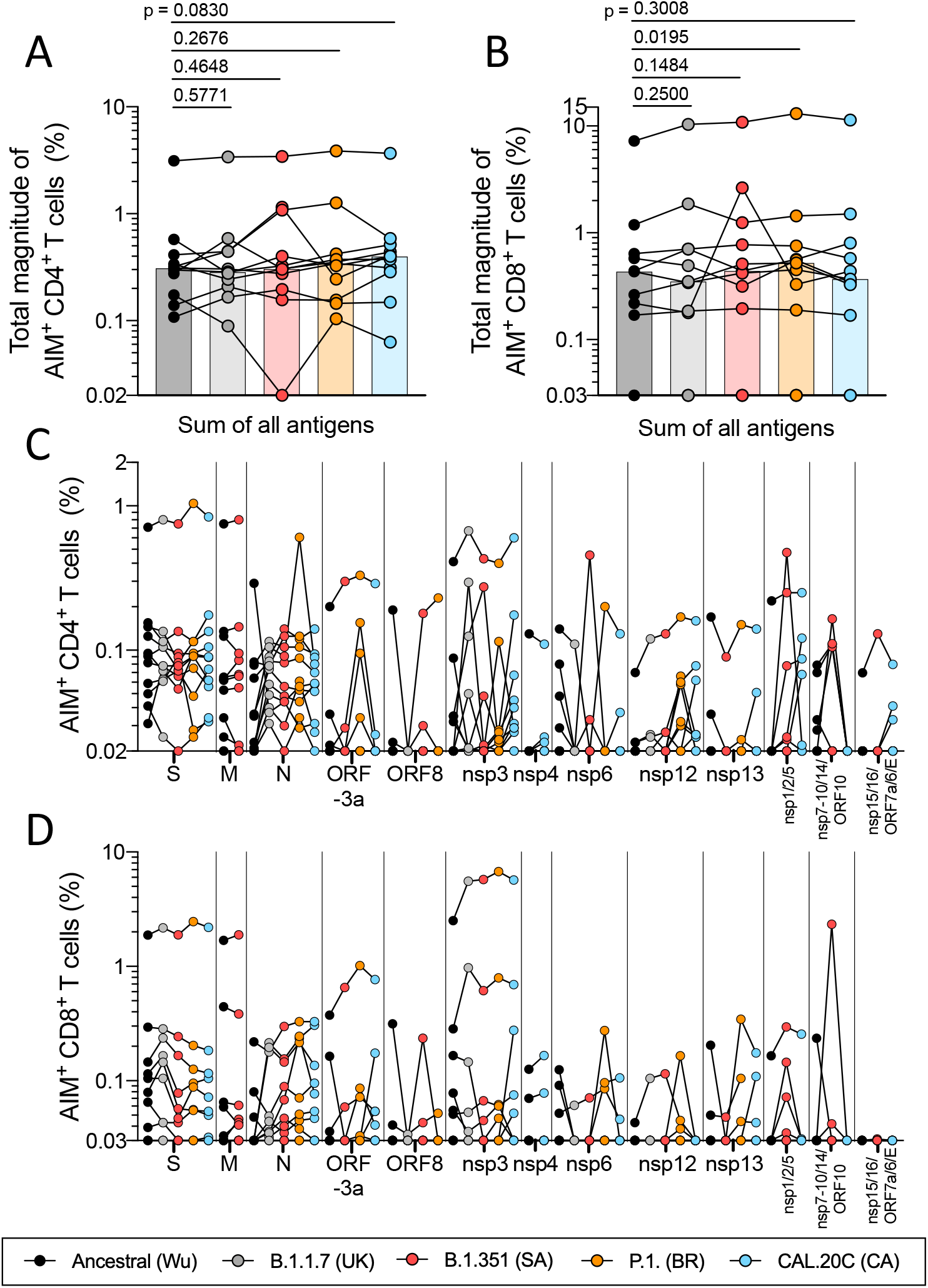
T cell responses of COVID-19 convalescent individuals against SARS-CoV-2 proteome for the different variants. PBMCs of the COVID-19 convalescent individuals (n=11) were stimulated with the MPs for the entire viral proteome corresponding to the ancestral reference strain (Wu, black) and the B.1.1.7 (UK, grey), B.1.351 (SA, red), P.1. (BR, orange) and CAL.20C (CA, light blue) SARS-CoV-2 variants. **A**) Percentages of AIM^+^ (OX40^+^CD137^+^) CD4^+^ T cells for the total reactivity. **B**) Percentages of AIM^+^ (CD69^+^CD137^+^) CD8^+^ T cells for the total reactivity. Bars represent the geometric mean. Paired comparisons of Wuhan versus each of the variants were performed by Wilcoxon tests. **C**) Percentages of AIM^+^ (OX40^+^CD137^+^) CD4^+^ T cells for each MP **D**) Percentages of AIM^+^ (OX40^+^CD137^+^) CD4+ T cells for each MP.

We previously showed that in COVID-19 convalescent subjects a set of 10 different antigens (nsp3, nsp4, nsp6, nsp12, nsp13, S, ORF3a, M, ORF8 and N) account for 83 and 81% of the total CD4^+^ and CD8^+^ T cell responses, respectively (Tarke et al., 2021). Here a similar overall pattern of dominant antigens was observed. When single proteins are considered, no variant pool showed a decrease in reactivity when a multi-hypothesis testing correction was applied (**Fig. 2C-D).** It is worth noting that this specific comparison is for illustration purposes only, as this study is not fully powered to rule out minor differences that could be observed in the individual antigens.

In conclusion, these experiments suggest that memory CD4^+^ or CD8^+^ T cells from individuals that have been infected with the ancestral SARS-CoV-2 strain recognize the ancestral reference strain and the variant genome-wide sequences with similar efficiency.

### CD4^+^ and CD8^+^ T cell antigenicity against Spike variant sequences in recent vaccines samples

We also studied T cell responses by individuals who received authorized mRNA COVID-19 vaccines. We focused our analysis on T cell reactivity to peptide pools spanning the Spike antigen of the ancestral strain, which is the basis of the presently used vaccines. For both CD4^+^ and CD8^+^ T cell reactivity, the magnitude of responses to pools encompassing the sequences from the ancestral Wuhan genome and the different variants considered range from a geomean of 0.15 to 0.19 for CD4^+^ T cells and a geomean of 0.16-0.24 for CD8^+^ T cells. Comparison of the variant pools to the ancestral sequence showed no significant difference for CD4^+^ T and CD8^+^ T cells reactivity, with the exception of the B.1.351 pools, where mild decreases of 29% and 33%, respectively, were observed (CD4: UK p=0.47; SA p=0.01; BR p=0.91; CA p=0.41; CD8: UK p=0.03; SA p=0.001; BR p=0.15 and CA p=0.02 by the Wilcoxon test) (**Fig. 3A-B**).

**Figure 3.**
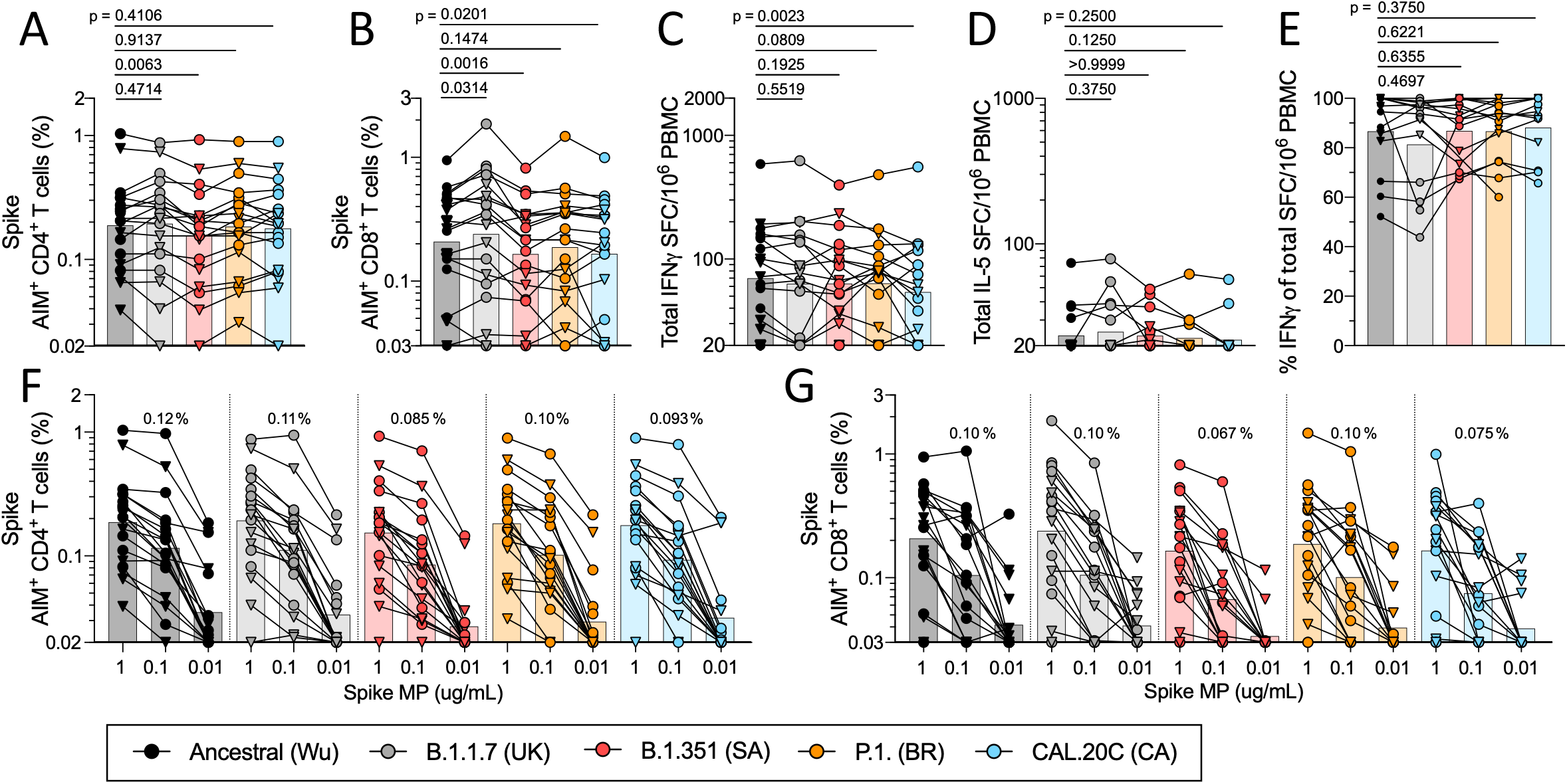
T cell responses of COVID-19 vaccinee individuals against SARS-CoV-2 Spike for the different variants. PBMCs of Pfizer/BioNTech BNT162b2 (n=8, triangles) and Moderna mRNA-1273 COVID-19 vaccines (n=11, circles) were stimulated with the Spike MPs corresponding to the ancestral reference strain (Wu, black) and the B.1.1.7 (UK, grey), B.1.351 (SA, red), P.1. (BR, orange) and CAL.20C (CA, light blue) SARS-CoV-2 variants. **A**) Percentages of AIM^+^ (OX40^+^CD137^+^) CD4^+^ T cells. **B**) Percentages of AIM^+^ (CD69^+^CD137^+^) CD8^+^ T cells. **C**) IFNγ spot forming cells (SFC) per million PBMCs **D**) IL-5 Spot forming cells (SFC) per million PBMCs. **E**) Percentages of IFNγ were calculated from the total IFNγ and IL-5 SFC per million PBMCs. Paired comparisons of the ancestral reference strain-based S MP versus each of the variants were performed by Wilcoxon test and are indicated by the p values in panels **A**-**E**. The data shown in panels **A** and **B** are also plotted showing the spike MPs titration (1 μg/mL, 0.1 μg/mL, 0.01 μg/mL) for CD4^+^ (**F**) and CD8^+^ (**G**) T cells in each SARS-CoV-2 variant. The geometric mean of the 0.1ug/mL condition is listed above each titration. In all panels, the bars represent the geometric mean.

The results from the FluoroSPOT assay system (**Fig. 3C-E**) showed good reactivity in terms of IFNγ, with geomean reactivity ranging from 54 to the 70 SFC per million PBMCs (**Fig. 3C**). Minimal IL-5 responses were observed, with geomean reactivity ranging from 22 to 25 SFC/10^6^, which is slightly above the limit of detection (**Fig. 3D**). On a per donor basis, the IFNγ response was found to account for more than 80% of the total response, on average (range 81% to 87%), irrespective of whether the ancestral strain or any of the variants was considered (**Fig. 3E**).

Similar to the experiments in convalescent donors, we also considered the dose response of the various Spike pools in terms of stimulation of CD4^+^ or CD8^+^ T cell specific responses for vaccinees. As shown in **Fig. 3F-G**, CD4^+^ and CD8^+^ T cell dose responses for the ancestral pools and the four variant pools were similar. Taken together these results indicate that the responses to the ancestral and variant Spike pools are similar for both CD4^+^ and CD8^+^ T cells in mRNA vaccinees.

### Conservation analysis of sets of defined CD4^+^ and CD8^+^ T cell epitopes

We recently reported a comprehensive study of epitopes recognized in convalescent subjects, leading to the identification of 280 different CD4^+^ T cell epitopes (Tarke et al., 2021). Here, we analyzed how many of those epitopes would be impacted by mutations in the different variants. As shown in **Fig. 4A**, we found that 89.6%, 90%, 94.3% and 97.1% (average 93%) of the CD4^+^ T cell epitopes identified by Tarke et al. are conserved in the B.1.1.7, B1.351, P.1 and the CAL20C variants. A similar pattern is observed when the magnitude of responses associated with the various epitopes is considered, rather than the simple number of epitopes (**Fig. 4B**). The fully conserved CD4^+^ T cell epitopes account for 84.4%, 88.1%, 95.7% and 97.8% (average 91.5%) of the total response, when comparing the B.1.1.7, B.1.351, P.1 and the CAL.20C variants, respectively.

**Figure 4.**
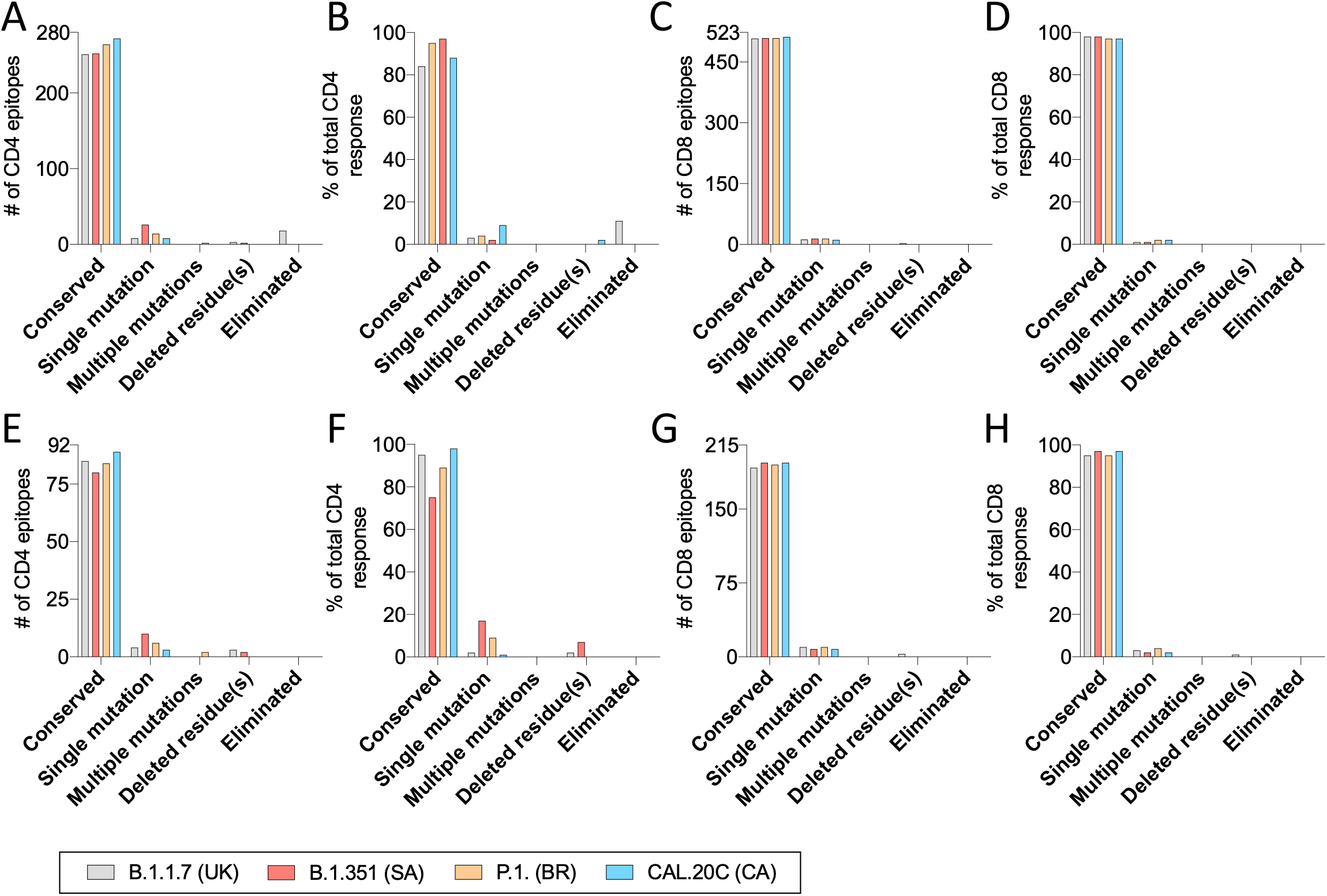
SARS-CoV-2 T cell epitope sequences affected by the variants. CD4^+^ and CD8^+^ T cell epitopes of the ancestral strain identified in a previous study (Tarke et al.) are analyzed as a function of the number and percentage of response that are or are not conserved across the B.1.1.7 (UK, grey), B.1.351 (SA, red), P.1. (BR, orange) and CAL.20C (CA, light blue) SARS-CoV-2 variants. The SARS-CoV-2 epitopes for the most immunodominant SARS-CoV-2 proteins in terms of numbers and percentage of response are shown for CD4^+^ (**A-B**) and CD8^+^ (**C-D**) T cells. The SARS-CoV-2 epitopes for the Spike protein only in terms of numbers and percentage of response are shown for CD4^+^ (**E-F**) and CD8^+^ (**G-H**) T cells.

The study of Tarke et al. also reported the identification of 523 CD8^+^ T cell epitope associated with unique HLA restrictions (Tarke et al., 2021). Performing a similar analysis as above, (**Fig. 4C**) we found that 508 (97.1%) of these 523 CD8^+^ T cell epitopes are totally conserved within the B.1.1.7 variant, 509 (97.3%) are conserved within the B.1.351 variant, 509 (97.3%) are conserved within the P.1 variant and 512 (97.9%) within the CAL.20C variant. Similarly, in terms of magnitude of CD8^+^ T cell responses associated with the various epitopes, totally conserved CD8^+^ T cell epitopes account for 98.3%, 98.4%, 97.9% and 97.8% of the total responses (**Fig. 4D**) for the B.1.1.7, B.1.351, P.1 and the CAL.20C variants respectively, with an average of 98.1%.

Finally, we analyzed the degree of CD4^+^ and CD8^+^ T cell epitope conservation if we restricted our analysis only to epitopes contained in the Spike antigen. The number of S-derived epitopes conserved at 100% sequence identity was, on average, 84.5% for the CD4^+^ T cell epitopes (**Fig. 4E),** and 95.3% for the CD8^+^ T cell epitopes (**Fig. 4G**). Similarly, in terms of magnitude of CD4^+^ T cell responses associated with the various S epitopes, totally conserved CD4^+^ T cell epitopes account for 95.5%, 75.3%, 89.8% and 98.3% of the total responses (**Fig. 4F**) for the B.1.1.7, B.1.351, P.1 and the CAL.20C variants respectively, with an average of 89.7%. In terms of the magnitude of CD8^+^ T cell responses, totally conserved epitopes account for 95.2%, 97.6%, 95.4% and 97.3% of the total responses (**Fig. 4H**) for the B.1.1.7, B.1.351, P.1 and the CAL.20C variants respectively, with an average of 96.4%.

While the restriction of the HLA class II epitopes in the Tarke et al. (Tarke et al., 2021) study was not unequivocally assigned, the restriction of the class I epitopes is implicitly inferred based on HLA allele specific predictions and testing in HLA matched donors. Accordingly, we further analyzed the extent to which the affected epitopes would be impacted by their respective associated mutation by determining, for each epitope/matching epitope variant, their predicted binding affinity for the corresponding putative HLA class I restriction element. Predicted binding capacity was determined using the NetMHCpan BA 4.1 tool provided by the IEDB’s analysis resource (Dhanda et al., 2019; Reynisson et al., 2020).

In the case of the B.1.1.7, B.1.351, P.1 and the CAL.20C variants, the % of mutations associated with no decrease in binding capacity, conservatively defined as a 2-fold reduction, was 73.3%, 78.6%, 78.6% and 45.5%, respectively (with the CAL.20.C variant having a smallest number of total mutations, noted above) (**Fig. S2** and **Table S3**). In conclusion, the analyses suggest that the vast majority of CD8^+^ T cell epitopes are unaffected by mutations found in all the different variants. The corresponding mutations are also predicted to have minor effects on the total T cell response, thus providing a molecular basis for the marginal impact on T cell reactivity by COVID-19 convalescent subjects and recipients of COVID-19 mRNA vaccines.

## DISCUSSION

The present study addresses a key knowledge gap pertaining to the potential of emergent SARS-CoV-2 variants to evade recognition by human immune responses. We focused on T cell responses elicited by either natural infection or vaccination with the Pfizer/BioNTech and Moderna COVID-19 mRNA vaccines. We found negligible effects on both CD4^+^ or CD8^+^ T cell responses to all four variants investigated, to include the B.1.1.7, B.1.351, P.1 and CAL.20C variants found in the UK, South Africa, Brazil and California, respectively. To more comprehensively assess T cell functionalities, the comparison between the original Wuhan isolate and the variants was performed utilizing different T cell methodologies, such as the AIM assay (quantifying T cells with a range of functionalities), and the FluoroSPOT assay (quantifying cells with specific cytokine-secreting activity). We also tested whether any of the variant sequences might be associated with an altered cytokine polarization; marginal IL-5 production was detected in any of the conditions tested. This is relevant, since it was reported that single amino acid replacements in an epitope sequence can lead to a change in the cytokines produced (Evavold and Allen, 1991; Sloan-Lancaster and Allen, 1996), and a Th2-like response pattern was initially hypothesized to be linked to adverse outcomes in SARS-specific responses (Peeples, 2020).

The data provide some positive news in light of justified concern over the impact of SARS-CoV-2 variants of concern on efforts to control and eliminate the present pandemic. Undoubtedly, several of the variants are associated with increased transmissibility, and also have been associated with decreased susceptibility to neutralizing antibodies from infected or vaccinated individuals. In contrast, the data presented here suggests that T cell responses are largely unaffected by the variants. While it is not anticipated that circulating memory T cells would be effective in preventing SARS-CoV-2 infection, it is plausible that they can reduce COVID-19 severity (Lipsitch et al., 2020; Sette and Crotty, 2021). Several lines of evidence support this notion, such as observations that early SARS-CoV-2 T cell responses are associated with milder COVID-19 (Rydyznski Moderbacher et al., 2020; Tan et al., 2021). Thus, the T cell response may contribute to limiting COVID-19 severity induced by variants that partially or largely escape neutralizing antibodies. This is consistent with T cell mediated immunity observed in humans against a different respiratory pathogen, influenza, for which heterologous immunity against diverse influenza strains is associated with memory T cells to conserved epitopes (Greenbaum et al., 2009; Sridhar et al., 2013; Wilkinson et al., 2012).

Our data also provide a molecular basis for the lack of impact of the mutations associated with the variants analyzed on T cell responses. Prior reports have identified a large number of T cell epitopes recognized throughout the SARS-CoV-2 proteome, including Spike (Ferretti et al., 2020; Keller et al., 2020; Le Bert et al., 2020; Nelde et al., 2020; Peng et al., 2020; Snyder et al., 2020). We furthered this point by an analysis of the Tarke et al. data set, showing that 93% of CD4^+^ T cell, and 97% of CD8^+^ T cell, epitopes are completely conserved in the variants. Further, we found that even in the epitopes affected by single mutations, no negative affect HLA binding capacity in the majority of cases is expected. The apparent higher conservation of CD8^+^ T cell epitopes is to be expected based on the shorter length of HLA class I binding peptides (usually 9-10 amino acids) as compared to their class II counterparts (13-17. This effect is counterbalanced by CD8^+^ T cells being generally less tolerant of amino substitutions as compared to CD4^+^ T cells (Grifoni et al., 2020a; Weiskopf et al., 2014). Overall, we observed that the effect of the variant mutations on the global CD4^+^ and CD8^+^ T cell responses was negligible.

Mutations associated with the variants could be reflective of adaptation in terms of optimizing replication or binding to ACE2, but also reflective of adaptation to escape immune recognition by antibodies (Andreano et al., 2020; Starr et al., 2020; Wang et al., 2021a; Wang et al., 2021b; Zahradník et al., 2021). Indeed, higher viral binding to a cellular receptor can be a mechanism of compensatory viral evolution in the presence of neutralizing antibodies (Hensley et al., 2009). In that respect, while mutation to escape antibody binding has been well documented for influenza (Andrews et al., 2015; Doud et al., 2018; Krammer et al., 2018) and SARS-CoV-2, immune escape at the level of T cell responses in human populations has not been reported for other acute respiratory infections. Because of HLA polymorphism, the epitope repertoire recognized is likely to be substantially different from one individual to the next, greatly decreasing the likelihood of immune escape by an acute virus. An advantage conferred to the virus by a mutation in a person would not be linked to an immune response escape advantage in a non-HLA matched individual. At the same time, our data does not rule out that each person could be strongly affected by the mutations of specific variants. For SARS-CoV-2, this property of T cell recognition is further enhanced by the fact that the T cell responses against SARS-CoV-2 are highly multi-antigenic and multi-specific, with tens of different epitopes recognized by CD4^+^ and CD8^+^ T cells in a given individual (Braun et al., 2020; Ferretti et al., 2020; Nelde et al., 2020; Tarke et al., 2021).

The results here have potential implications for engineering coronavirus vaccines with broader protective immunity against variants of concern. Clearly the most straightforward path is to update the current vaccines to target a variant Spike, given how highly successful several COVID-19 vaccines have proven to be against the parental SARS-CoV-2 strain. Our results suggest that a parallel alternative approach could involve inclusion of additional antigens and epitopes, perhaps selected on the basis of low mutational propensity (Gaiha et al., 2019), to ensure that neutralizing antibodies are complemented with T cell responses to minimize COVID-19 morbidity and mortality.

### Limitations and future directions

The present study did not assess decreases in antibody reactivity, as several other studies have already investigated this matter (Edara et al., 2021; Greaney et al., 2021; Muik et al., 2021; Shen et al., 2021; Skelly et al., 2020; Stamatatos et al., 2021; Supasa et al., 2021; Wang et al., 2021a; Wang et al., 2021b; Wibmer et al., 2021; Wu et al., 2021). Further, our studies utilized overlapping peptide pools, and as such we could not exclude that some of the mutations might involve alterations in terms of antigen processing for either class I or class II, which would be undetected by using pools of “preprocessed” peptides. The number of donors studied was also limited, although variability in T cell reactivity suggestive of large variant-associated effects were not observed. While we have no reason to suspect that substantial differences might exist between the epitope specificity of responses elicited by different vaccines, our study did not address this point. Our study was designed to test for differences in overall response to the different variants and was not powered or designed to investigate differences between the mRNA vaccines. Furthermore, in our study we tested samples from convalescent donors infected before October 2020; thus, it is unlikely that any of the donors would have been infected by any of the variants, as this date precedes their diffusion to appreciable degree in the US and California. Finally, we have only investigated whether responses induced by the ancestral reference sequence are able to cross-recognize variant sequences, as this is relevant to the current situation. We have not examined whether responses induced by an infection with a variant sequence will be able to cross-recognize the ancestral reference sequence present in the currently approved vaccines.

## ACKNOWLEDGEMENTS

This study has been funded by the NIH NIAID (award AI142742 to S.C., A.S., contract Nr. 75N9301900065 to A.S. and D.W., contract Nr. 75N93019C00001 to A.S. and B.P., NIH grant U01 CA260541-01 to DW, K08 award AI135078 to J.D., and AI036214 to D.S. and HHSN75N93019C00076 to R.H.S). Additional support has been provided by UCSD T32s (AI007036 and AI007384 to S.A.R and S.I.R) and the Jonathan and Mary Tu Foundation (D.S.). A.T. was supported by a PhD student fellowship through the Clinical and Experimental Immunology Course at the University of Genoa, Italy. We thank Gina Levi and the LJI clinical core for assistance in sample coordination and blood processing. We gratefully thank the authors from the originating laboratories responsible for obtaining the specimens, as well as the submitting laboratories where the genome data were generated and shared via GISAID, and on which this research is based. We would like to thank Vamseedhar Rayaprolu and Erica Ollmann Saphire for providing the recombinant SARS-CoV-2 Receptor Binding Domain (RBD) protein used in the ELISA assay.

## AUTHOR CONTRIBUTIONS

Conceptualization: A.T., A.G., S.C. and A.S.; Data curation and bioinformatic analysis, Y.Z. R.H.S. B.P.; Formal analysis: A.T., J.S., A.G.; Funding acquisition: S.C., A.S., D.W., S.I.R., S.A.R., and J.M.D.; Investigation: A.T., N.M., A.Su., B.G. J.S., D.W. A.S and A.G.; Project administration: A.F. Resources: S.I.R., S.A.R., J.D.; Supervision: J.S., S.C., D.W., A.S., R.d.S. and A.G.; Writing: A.T., D.W., S.C., A.S., and A.G.

## SUPPLEMENTAL MATERIALS

### SUPPLEMENTAL FIGURE LEGENDS

**Figure S1.**
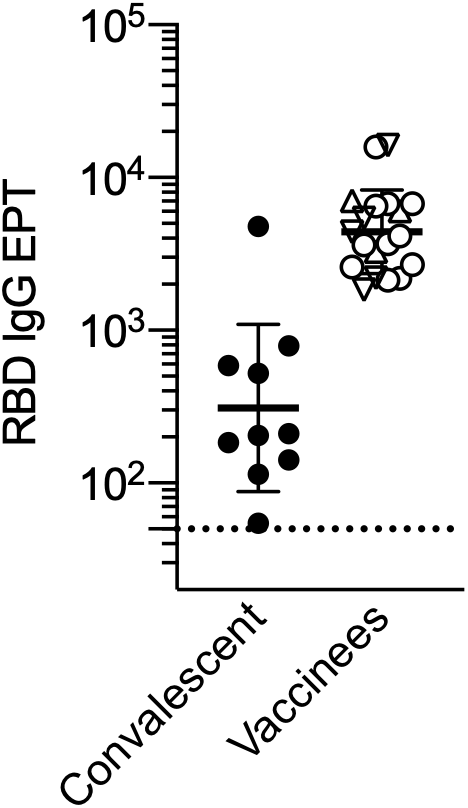
SARS-COV-2 serology of the all the cohorts analyzed in this study. Related to Figures 1, 2 and 3 and Table 1. Spike RBD serology in COVID-19 convalescents (n=11) and COVID-19 vaccines (n=19).

**Figure S2.**
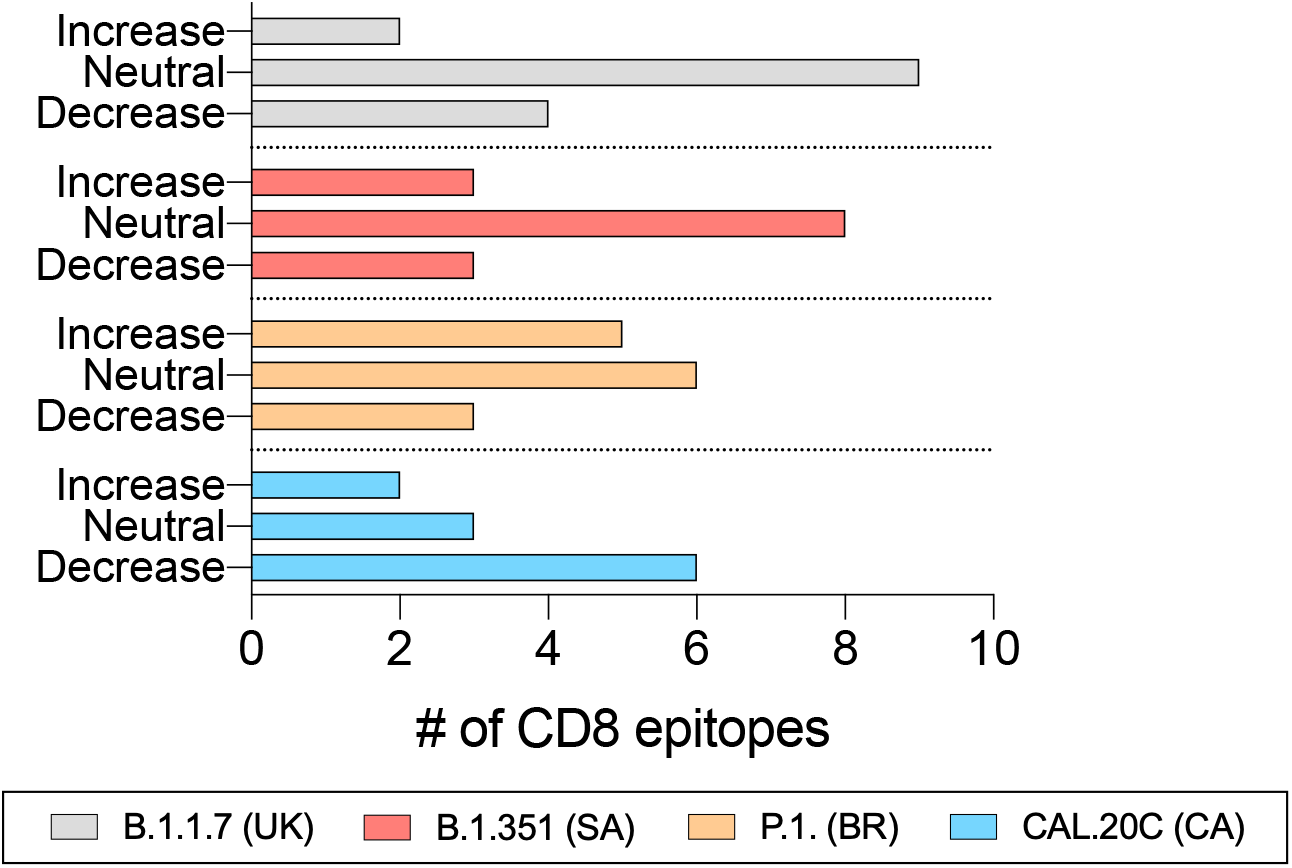
Effect of mutations on CD8 epitope predicted HLA class I binding capacity. Related to Figure 4. For each CD8^+^ T cell epitope associated with a mutation found in the respective variants, the predicted HLA binding capacity of original sequence and the mutated sequence was calculated. Based on the results, each instance was categorized as a function of whether the binding capability of the mutated peptide is increased (>2-fold), neutral or decreased (<2-fold). Each analysis is done separately for the B.1.1.7 (UK, grey), B.1.351 (SA, red), P.1. (BR, orange) and CAL.20C (CA, light blue) SARS-CoV-2 variants.

**Figure S3.**
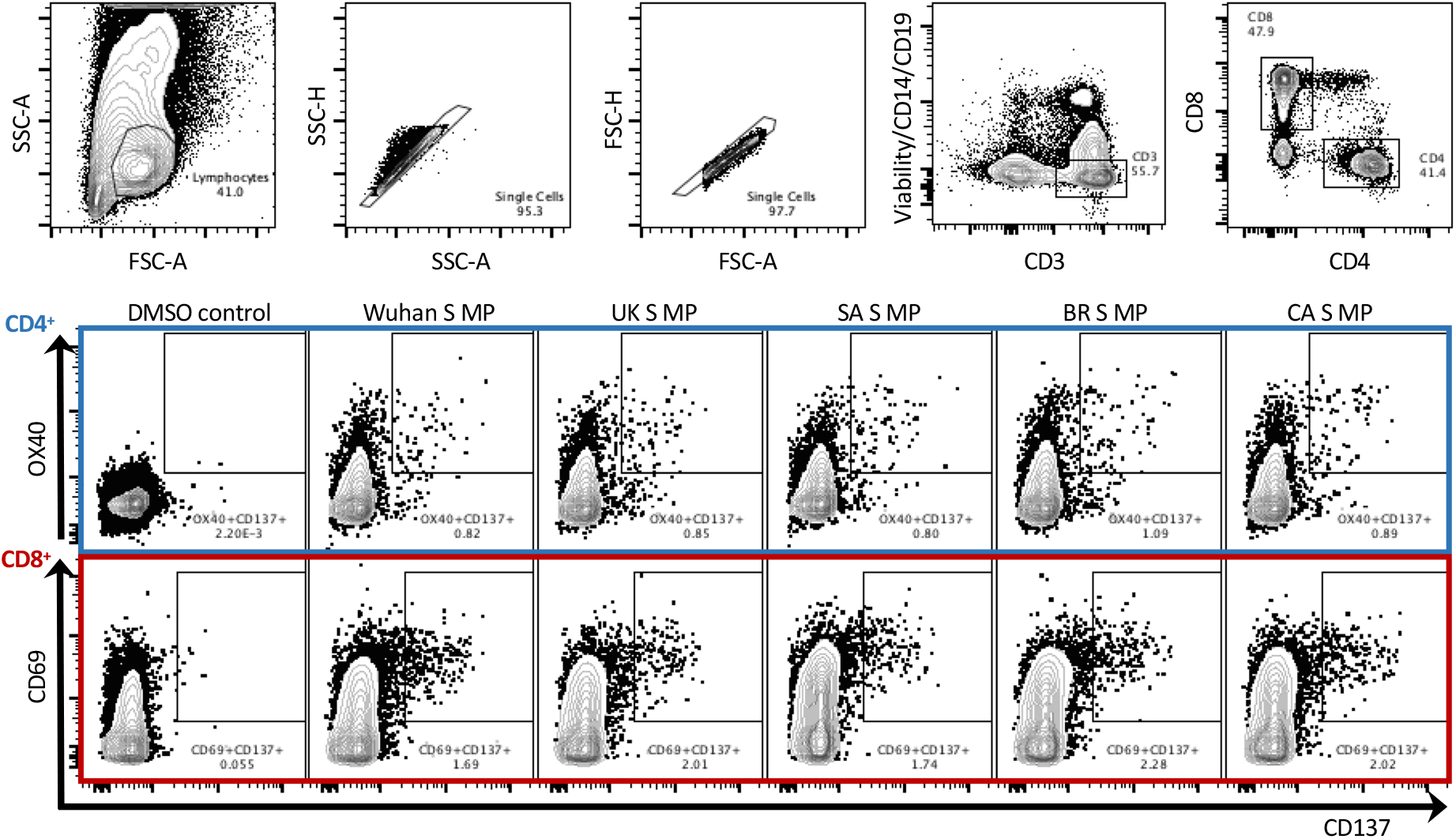
Gating strategy. Related to Figures 1, 2, 3 and 4. Representative graphs illustrating the gating strategy used in the flow cytometry AIM assays in order to define antigen-specific CD4^+^ (outlined in blue) and CD8^+^ (outlined in red) T cells by the expression of OX40^+^CD137^+^ and CD69^+^ CD137^+^, respectively. These graphs depict one of the COVID-19 convalescent donors from this study and are representative of the gating strategy utilized with all donors tested.

### SUPPLEMENTAL TABLE LEGENDS

**Table S1.** Related to Figures 1, 2, 3 and 4. List of amino acid positions and relative amino acid changes in the different variants studied with respect to the ancestral Wuhan strain.

| Protein | Amino acid position | Ancestral (Wu) | B.1.1.7 (UK) | B.1.351 (SA) | P.1. (BR) | CAL.20C (CA) |
| --- | --- | --- | --- | --- | --- | --- |
| S | 13 | S |  |  |  | I |
| S | 18 | L |  | F | F |  |
| S | 20 | T |  |  | N |  |
| S | 26 | P |  |  | S |  |
| S | 69 | H | Del |  |  |  |
| S | 70 | V | Del |  |  |  |
| S | 80 | D |  | A |  |  |
| S | 138 | D |  |  | Y |  |
| S | 145 | Y | Del |  |  |  |
| S | 152 | W |  |  |  | C |
| S | 190 | R |  |  | S |  |
| S | 215 | D |  | G/H |  |  |
| S | 241 | L |  | Del |  |  |
| S | 242 | L |  | Del |  |  |
| S | 243 | A |  | Del |  |  |
| S | 417 | K |  | N | T |  |
| S | 452 | L |  |  |  | R |
| S | 484 | E |  | K | K |  |
| S | 501 | N | Y | Y | Y |  |
| S | 570 | A | D |  |  |  |
| S | 614 | D | G | G | G | G |
| S | 655 | H |  |  | Y |  |
| S | 681 | P | H |  |  |  |
| S | 701 | A |  | V |  |  |
| S | 716 | T | I |  |  |  |
| S | 938 | L |  |  |  | F |
| S | 982 | S | A |  |  |  |
| S | 1027 | T |  |  | I |  |
| S | 1118 | D | H |  |  |  |
| S | 1176 | V |  |  | F |  |
| S | 1191 | K |  |  |  | N |
| M | 162 | K |  | N |  |  |
| N | 3 | D | L |  |  |  |
| N | 13 | P |  | S |  |  |
| N | 32 | R |  | H |  |  |
| N | 80 | P |  |  | R |  |
| N | 203 | R | K |  | K | K |
| N | 204 | G | R |  | R | R |
| N | 205 | T |  | I |  | I |
| N | 212 | G |  | C |  |  |
| N | 234 | M |  |  |  | I |
| N | 235 | S | F |  |  |  |
| E | 71 | P |  | L |  |  |
| ORF3a | 57 | Q |  | H |  | H |
| ORF3a | 131 | W |  | L |  |  |
| ORF3a | 171 | S |  | L |  |  |
| ORF3a | 253 | S |  |  | P |  |
| ORF7a | 93 | V |  | F |  |  |
| ORF8 | 27 | - | Stop |  |  |  |
| ORF8 | 92 | E |  |  | K |  |
| ORF8 | 121 | I |  | L |  |  |
| nsp1 | 109 | P |  | S |  |  |
| nsp2 | 85 | T |  | I |  | I |
| nsp2 | 339 | G |  | S |  |  |
| nsp2 | 366 | S |  | T |  |  |
| nsp2 | 427 | Q |  | H |  |  |
| nsp2 | 563 | E |  | D |  |  |
| nsp3 | 183 | T | I |  |  |  |
| nsp3 | 186 | T |  |  | A |  |
| nsp3 | 370 | S |  |  | L |  |
| nsp3 | 778 | P |  |  |  | S |
| nsp3 | 837 | K |  | N |  |  |
| nsp3 | 890 | A | D |  |  |  |
| nsp3 | 926 | C |  | S |  |  |
| nsp3 | 977 | K |  |  | Q |  |
| nsp3 | 1180 | T |  | I |  |  |
| nsp3 | 1412 | I | T |  |  |  |
| nsp3 | 1778 | N |  | S |  |  |
| nsp4 | 395 | S |  |  |  | T |
| nsp5 | 90 | K |  | R |  |  |
| nsp5 | 193 | A |  | V |  |  |
| nsp6 | 106 | S | Del | Del | Del |  |
| nsp6 | 107 | G | Del | Del | Del |  |
| nsp6 | 108 | F | Del | Del | Del |  |
| nsp6 | 125 | L |  |  |  | F |
| nsp6 | 135 | G |  | S |  |  |
| nsp6 | 149 | V |  | F |  |  |
| nsp6 | 167 | L |  |  |  | F |
| nsp9 | 65 | I |  |  |  | V |
| nsp10 | 105 | N |  | K |  |  |
| nsp12 | 323 | P | L | L | L | L |
| nsp13 | 53 | P |  |  |  | L |
| nsp13 | 209 | V |  |  |  | F |
| nsp13 | 260 | D |  |  |  | Y |
| nsp13 | 341 | E |  |  | D |  |
| nsp13 | 588 | T |  | I |  |  |
| nsp14 | 177 | L |  | F |  |  |
| nsp14 | 326 | F |  |  |  | L |
| nsp14 | 328 | V |  | F |  |  |
| nsp15 | 91 | D |  |  |  | Y |

**Table S2.**
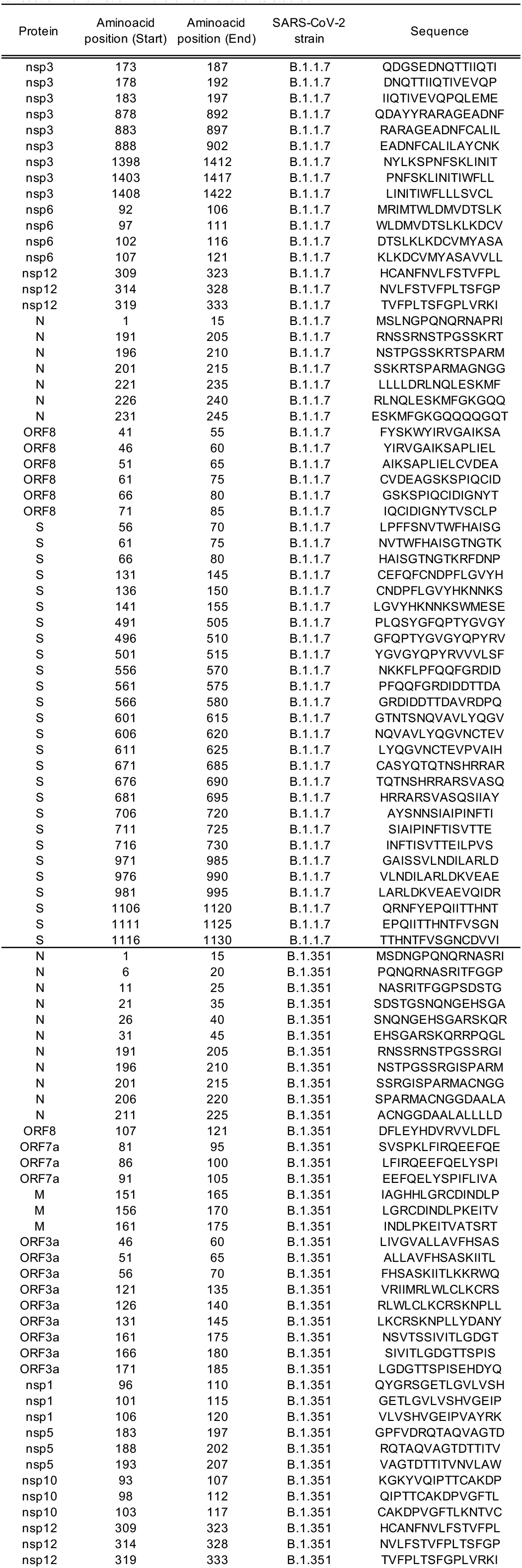
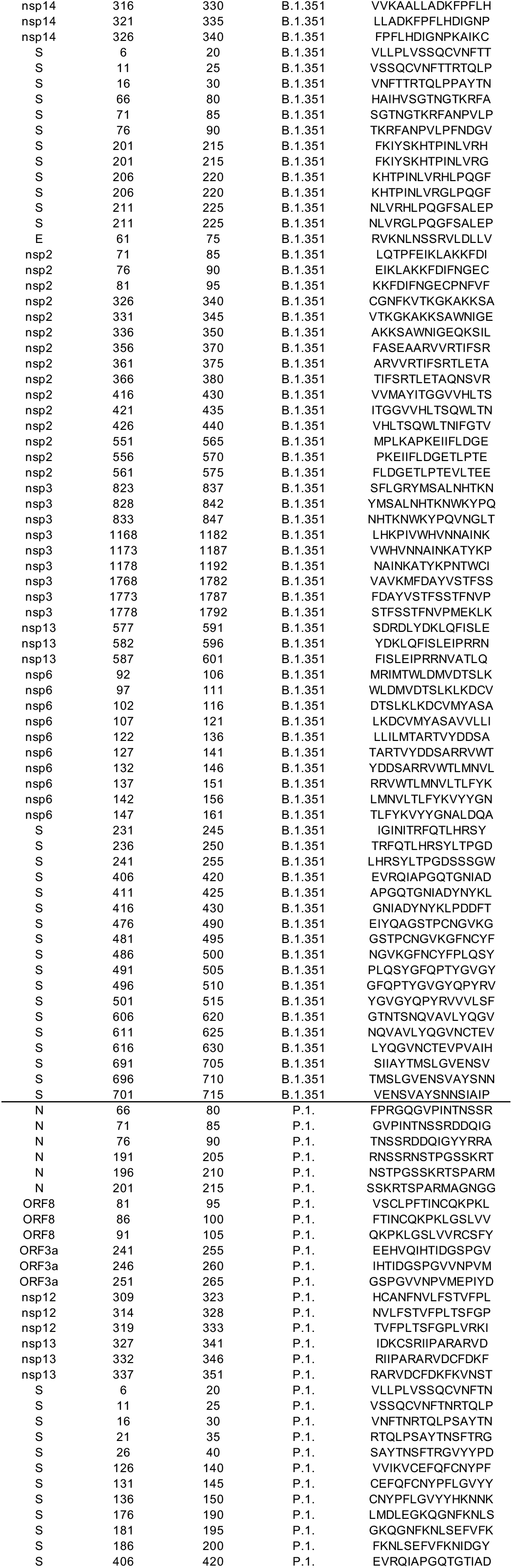
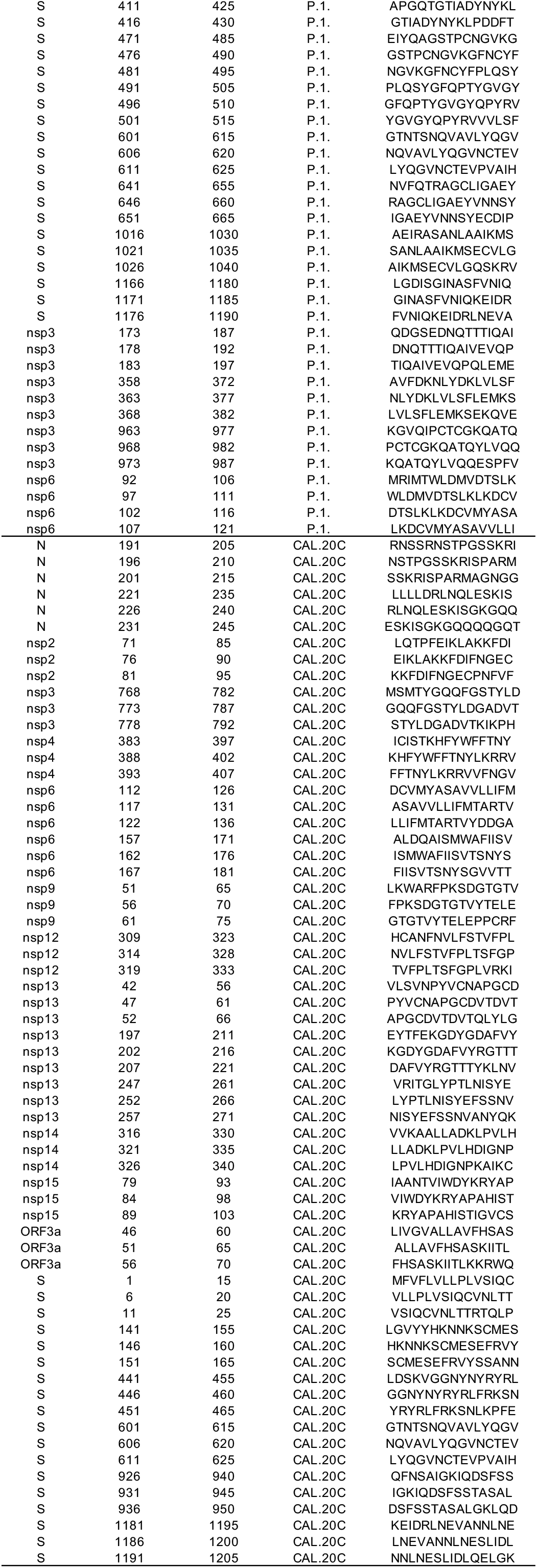
Related to Figures 1, 2, 3 and 4. List of mutated peptides with respect to the ancestral Wuhan strain in the different variants studied.

**Table S3.**
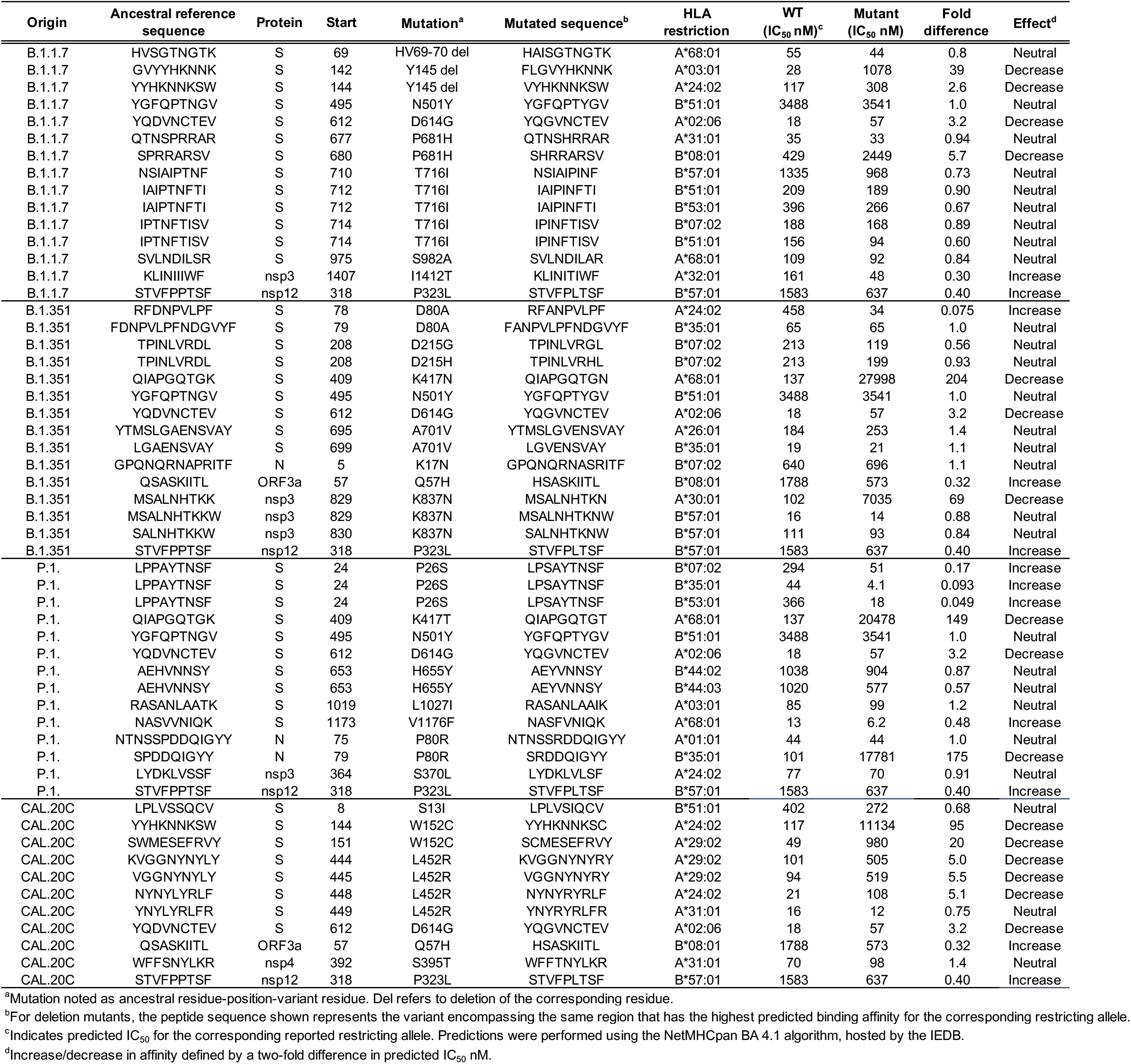
Related to Figure 4. Effect of mutations on CD8 epitope HLA class I binding capacity.

## STAR METHODS

### RESOURCE AVAILABILITY

#### Lead Contact

Further information and requests for resources and reagents should be directed to and will be fulfilled by the Lead Contact, Dr. Alessandro Sette.

#### Materials Availability

Aliquots of synthesized sets of peptides utilized in this study will be made available upon request. There are restrictions to the availability of the peptide reagents due to cost and limited quantity.

#### Data and Code Availability

The published article includes all data generated or analyzed during this study, and summarized in the accompanying tables, figures and supplemental materials.

## EXPERIMENTAL MODEL AND SUBJECT DETAILS

### Human Subjects

#### Convalescent COVID-19 Donors

Convalescent donors were enrolled at either a UC San Diego Health clinic under the approved IRB protocols of the University of California, San Diego (UCSD; 200236X), or at the La Jolla Institute (LJI; VD-214). All donors were California residents and samples were collected from August to October 2020, before any of the SARS-CoV-2 variants described herein had been detected in California. These donors were referred to the study by a health care provider or were self-referred. The CRO BioIVT provided additional cohorts of COVID-19 convalescent donors who had been confirmed positive for COVID-19 by PCR following the resolution of symptoms. The total cohort of convalescent donors represented both sexes (27% male, 73% female) and ranged from 21 to 57 years of age (median 39 years). All samples were confirmed seropositive against SARS-CoV-2 by ELISA, as described below. Details of this convalescent COVID-19 cohort are listed in Table 1. All convalescent COVID-19 donors provided informed consent to participate in the present and future studies at the time of enrollment.

#### COVID-19 vaccinees

The La Jolla Institute recruited 19 healthy adults who had received the first and second dose of the Pfizer/BioNTech BNT162b2 (n=8) or Moderna mRNA-1273 COVID-19 vaccines (n=11). Blood draws took place under IRB approved protocols two to four weeks after the second dose of the vaccine was administered. All donors had their SARS-CoV-2 antibody titers measured by ELISA, as described below. The cohort of vaccines represented ranged from 22 to 67 years of age (median 43 years) and represented both sexes (26% male, 74% female). At the time of enrollment in the study, all donors gave informed consent.

## METHOD DETAILS

### Isolation of peripheral blood mononuclear cells (PBMCs) and plasma

Collection and processing of blood samples was performed as previously described (Dan et al., 2021; Tarke et al., 2021). Briefly, whole blood was collected in heparin coated blood bags or in ACD tubes and centrifuged for 15 minutes at 1850 rpm to separate the cellular fraction from the plasma. The plasma was then removed and stored at −20°C. The cellular fraction next underwent density-gradient sedimentation using Ficoll-Paque (Lymphoprep, Nycomed Pharma, Oslo, Norway) to separate the PBMCs as previously described (Weiskopf 2013). Isolated PBMCs were cryopreserved in cell recovery media containing 10% DMSO (Gibco), supplemented with 90% heat inactivated fetal bovine serum (FBS; Hyclone Laboratories, Logan UT) and stored in liquid nitrogen until used in the assays.

### SARS-CoV-2 RBD ELISA

Serology to SARS-CoV-2 was determined for all donor cohorts as previously described (Rydyznski Moderbacher et al., 2020). Briefly, 96-well half-area plates (ThermoFisher 3690) were coated with 1 ug/mL SARS-CoV-2 Spike (S) Receptor Binding Domain (RBD) and incubated at 4°C overnight. The next day plates were blocked at room temperature for 2 hours with 3% milk in phosphate buffered saline (PBS) containing 0.05% Tween-20. Heat-inactivated plasma was added to the plates for an additional 90-minute incubation at room temperature followed by incubation with the conjugated secondary antibody, detection, and subsequent data analysis by reading the plates on Spectramax Plate Reader at 450 nm using the SoftMax Pro. The limit of detection (LOD) was defined as 1:3. Limit of sensitivity (LOS) for SARS-CoV-2 infected individuals was established based on uninfected subjects, using plasma from normal healthy donors not exposed to SARS-CoV-2.

### Mutation analysis of SARS-CoV-2 UK, California, South Africa and Brazil variants

Genome sequences for the variant viruses were downloaded from GISAID. These sequences were screened to select those without ambiguous residues and generated from Illumina sequencing technologies using an in-house sequence QC script. The selected genomic sequences were then translated into protein amino acid sequences using the VIGOR4 tool available on the Virus Pathogen Resource (ViPR)(Pickett et al., 2012). Sequence variations in the variant viruses were derived by comparison with Wuhan-1 (NC_045512.2). One or more representative sequences were considered for the UK (EPI_ISL_601443), Brazilian (EPI_ISL_804823), Californian (EPI_ISL_847619; EPI_ISL_847621; EPI_ISL_847643) and South Africa (EPI_ISL_660629; EPI_ISL_736930; EPI_ISL_736932; EPI_ISL_736944; EPI_ISL_736966; EPI_ISL_736971; EPI_ISL_736973; EPI_ISL_825104; EPI_ISL_825120; EPI_ISL_825131) variants. A summary of all the amino acids mutated in the different variants respect to the Wuhan sequence and considered in this study is available in **Table S1**.

### SARS-CoV-2 Wuhan and variant peptide synthesis and pooling

Peptides were synthesized that spanned entire SARS-CoV-2 proteins and corresponded to the ancestral Wuhan sequence or the B.1.1.7 (UK), B.1.351 (SA), P.1 (BR) and CAL.20C (CA) SARS-CoV-2 variants. Peptides were 15-mers overlapping by 10 amino acids and were synthesized as crude material (TC Peptide Lab, San Diego, CA). All peptides were individually resuspended in dimethyl sulfoxide (DMSO) at a concentration of 10–20 mg/mL. Megapools (MP) for each antigen were created by pooling aliquots of these individual peptides, undergoing another lyophilization, and resuspending in DMSO at 1 mg/mL.

### Bioinformatic analysis of T cell epitopes

The binding capacity of SARS-CoV-2 T cell epitopes, and their corresponding variant-derived peptides, for their putative HLA class I restricting allele(s) was determined utilizing the NetMHCpan BA 4.1 algorithm (Reynisson et al., 2020), as implemented by the IEDB’s analysis resource (Dhanda et al., 2019; Vita et al., 2019). Predicted binding is expressed in terms of IC_50_ nM. For each epitope-variant pair a ratio of affinities (WT/variant) was determined. Ratios >2, indicating a 2-fold or greater increase in affinity due to the mutation, were categorized as an increase in binding capacity, and <0.5 as a decrease; ratios between 0.5 and 2 were designated as neutral.

### Flow cytometry-based AIM assay

Activation induced cell marker (AIM) assay has previously been described in detail elsewhere (da Silva Antunes et al., 2021; Dan et al., 2021; Reiss et al., 2017). In summary, PBMCs were cultured for in the presence of SARS-CoV-2 specific (Wuhan or variant) MPs [1 μg/ml] in 96-well U-bottom plates at a concentration of 1×10^6^ PBMC per well. As a negative control, an equimolar amount of DMSO was used to stimulate the cells in triplicate wells and as positive controls phytohemagglutinin (PHA, Roche, 1μg/ml) and a cytomegalovirus MP (CMV, combining CD4 and CD8 MPs, 1μg/ml) were also included. After incubation for 20–24 hours at 37°C, 5% CO_2_, the cells were stained with CD3 BUV805 or CD3 AF700 (4:100 or 4:100; BD Biosciences Cat# 612895 or Life Technologies Cat# 56-0038-42, respectively), CD4 BV605 (4:100; BD Biosciences Cat# 562658), CD8 BUV496 or BV650 (2:100 or 4:100; BD Biosciences Cat# 612942 or Biolegend Cat# 301042), and Live/Dead eFluor506 (5:1000; eBioscience Cat# 65-0866-14). Cells were also stained to measure activation with the following markers: CD137 APC (4:100; Biolegend Cat# 309810), OX40 PE-Cy7 (2:100; Biolegend Cat#350012), and CD69 PE (10:100; BD Biosciences Cat# 555531). All samples were acquired on a ZE5 5-laser or 4-laser cell analyzer (Bio-rad laboratories) and analyzed with FlowJo software (Tree Star). In the resulting data generated from the AIM assays, the background was removed from the data by subtracting the average of the % of AIM^+^ cells plated in triplicate wells stimulated with DMSO. The Stimulation Index (SI) was calculated by dividing the % of AIM^+^ cells after SARS-CoV-2 stimulation with the average % of AIM^+^ cells in the negative DMSO control. An SI greater than 2 and a minimum of 0.02 % or 0.03 % AIM^+^ CD4^+^ or CD8^+^ cells, respectively, after background subtraction was considered to be a positive response. The gates for AIM^+^ cells were drawn relative to the negative and positive controls for each donor. A representative example of the gating strategy is depicted in **Fig. S3**.

### FluoroSPOT assays

96-well FluoroSpot plates were coated with anti-cytokine antibodies for IFNγ and IL-5 (mAbs 1-D1K and TRFK5, respectively; Mabtech, Stockholm, Sweden) at a concentration of 10μg/mL. PBMCs were stimulated in triplicate at a density of 200×10^3^ cells/well with S MPs corresponding to each of the SARS-CoV-2 variants analyzed (1μg/mL), PHA (1μg/mL), and DMSO (0.1%), as positive and negative controls respectively. After 20 hours of incubation at 37°C, 5% CO_2_, cells were discarded and plates were washed before the addition of cytokine antibodies (mAbs 7-B6-1-BAM and 5A10-WASP; Mabtech, Stockholm, Sweden). After a 2-hour incubation, plates were washed again with PBS/0.05% Tween20 and incubated for 1 hour with fluorophore-conjugated antibodies (Anti-BAM-490 and Anti-WASP-640). An AID iSPOT FluoroSpot reader (AIS-diagnostika, Germany) was used to count the fluorescent spots that resulted from cells secreting IFNγ and IL-5. Each peptide MP was considered positive compared to the DMSO negative control based on the following criteria: 20 or more spot forming cells (SFC) per 10^6^ PBMC after subtraction, a stimulation index (S.I.) greater than 2, and a p value <0.05 by either a Poisson or T test calculated between the triplicates of the MP and the relative negative control.

## QUANTIFICATION AND STATISTICAL ANALYSIS

Data and statistical analyses were performed in FlowJo 10 and GraphPad Prism 8.4, unless otherwise stated. Statistical details of the experiments are provided in the respective figure legends and in each method section pertaining the specific technique applied. Data plotted in logarithmic scales are expressed as geometric mean. Statistical analyses were performed using Wilcoxon matched-pairs signed rank test for paired comparisons. Multi-hypothesis testing corrections (MHTC) have not been applied in the study by design. The study is not designed or powered to address differences across different proteins. The primary hypothesis is that no significant differences are observed across the different variants, and this is more stringently addressed avoiding to correct for MHTC, since a difference that is not significant would remain so even after corrections. Therefore, reporting the data without applying MHTC is a more stringent criterion which is appropriately being applied in this case to avoid false negatives. Details pertaining to significance are also noted in the respective figure legends.

## DECLARATION OF INTEREST

A.S. is a consultant for Gritstone, Flow Pharma, Oxford Immunetech, Caprion, Arcturus and Avalia. S.C. is a consultant for Avalia. All other authors declare no conflict of interest. LJI has filed for patent protection for various aspects of vaccine design and identification of specific epitopes.

